# An integrated *in silico* immuno-genetic analytical platform provides insights into COVID-19 serological and vaccine targets

**DOI:** 10.1101/2020.05.11.089409

**Authors:** Daniel Ward, Matthew Higgins, Jody E. Phelan, Martin L. Hibberd, Susana Campino, Taane G Clark

## Abstract

**Background:** The COVID-19 pandemic, caused by the SARS-CoV-2 virus, has a major global health and socio-economic burden. It has instigated the mobilisation of resources into the development of control tools, such as diagnostics and vaccines. The poor performance of some diagnostic serological tools has emphasised the need for up to date immune-informatic analyses to inform the selection of viable targets for further study. This requires the integration and analysis of genetic and immunological data for SARS-CoV-2 and its homology with other human coronavirus species to understand cross-reactivity.

**Methods:** We have developed an online “immuno-analytics” resource to facilitate SARS-CoV-2 research, combining an extensive B/T-cell epitope mapping and prediction meta-analysis, and human CoV sequence homology mapping and protein database annotation, with an updated variant database and geospatial tracking for >7,800 non-synonymous mutation positions derived from >150,000 whole genome sequences. To demonstrate its utility, we present an integrated analysis of SARS-CoV-2 spike and nucleocapsid proteins, both being vaccine and serological diagnostic targets, including an analysis of changes in relevant mutation frequencies over time.

**Results:** Our analysis reveals that the nucleocapsid protein in its native form appears to be a sub-optimal target for use in serological diagnostic platforms. The most frequent mutations were the spike protein D614G and nsp12 L314P, which were common (>86%) across all the geographical regions. Some mutations in the spike protein (e.g. A222V and L18F) have increased in frequency in Europe during the latter half of 2020, detected using our automated algorithms. The tool also suggests that orf3a proteins may be a suitable alternative target for diagnostic serologic assays in a post-vaccine surveillance setting.

**Conclusions:** The immuno-analytics tool can be accessed online (http://genomics.lshtm.ac.uk/immuno) and will serve as a useful resource for biological discovery and surveillance in the fight against SARS-CoV-2. Further, the tool may be adapted to inform on biological targets in future outbreaks, including potential emerging human coronaviruses that spill over from animal hosts.

## Background

COVID-19, the disease caused by the SARS-CoV-2 virus, was first characterised in the city of Wuhan, Hubei, and has now spread to 190 countries. With over 60 million confirmed cases worldwide and more than 1.26 million deaths, the COVID-19 pandemic has placed a high burden on the world’s healthcare infrastructure and economies, with projected final costs of 28 trillion or 31% of the global gross domestic product [1,2]. The majority of infections are either asymptomatic or result in mild flu-like disease, with severe cases of viral pneumonia affecting between 1.0% (≥20 years) and 18.4% (≥80 years) of diagnosed patients [3]. Its variable infection outcome, mode of transmission and incubation period together have enhanced the ability of the pathogen to spread efficiently worldwide. As a result, there has been an urgent push for the development of diagnostics, therapeutics and vaccines to aid control efforts.

Current front-line diagnostic strategies apply a quantitative reverse transcription PCR (RT-qPCR) assay on patient nasopharyngeal swabs, using primer/probe sets targeting the *nsp10, RdRp, nsp14*, envelope and nucleocapsid genes; tests endorsed by a number of agencies and health systems [4,5] Patients hospitalised with severe respiratory disease who are RT-qPCR negative may be diagnosed radiographically (chest x-ray or computerised tomography scan), but in resource-poor and high infection rate settings these methods maybe unviable. Considering the inherent limitations in the sample collection process and transient viral load, RNA detection-based diagnostics can vary in their sensitivity. The demand for serological diagnostics is high, particularly because these tests are capable of detecting SARS-CoV-2 antibodies, a biomarker indicative of infection, present after the infection has been cleared by the immune system. These tools are essential to address crucial questions like how many people have been infected within a population, including those who may have been asymptomatic, and how long immunity can last post infection.

Numerous lateral flow rapid diagnostic tests (RDTs) and enzyme-linked immunosorbent assays (ELISA) tests have been developed, including an approved IgM/IgG RDT which uses the nucleocapsid protein as a target for the detection of seroconverted individuals [6]. Other assays use the spike protein as an antigen, with some using the receptor binding domain (RBD) as a target, a region with a high level of diversity between alphacoronavirus species [7]. Unlike RNA detection methods, these platforms can identify convalescent patients, which further informs outbreak control efforts. Long-term control strategies will involve vaccine roll-out. As of November 2020, there were more than 50 vaccines at development phase 1 or greater, with at least 10 vaccines in phase 3 [8,9]. Vaccines at the forefront, include those based on a non-replicating adenovirus vector base (ChAdOx-nCoV-19 and Ad5-nCov), an LNP-encapsulated mRNA (BNT162 and mRNA-1273), protein subunit (NVX-CoV2373) or inactivated virus (BBIBP-CorV and CoronaVac) [8,9].

The discovery, development and management of efficacious vaccines and sensitive and specific serological diagnostics are both dependant on the availability of up-to-date information on viral evolution and immune-informatic analyses. The identification of variable or conserved regions in the proteome of SARS-CoV-2 can inform the rational selection of reverse-design targets in both vaccinology and diagnostic fields, as well as indicate immunologically relevant regions of interest for further studies to characterise SARS-CoV-2 immune responses. Whilst the availability of biological data for SARS-CoV-2 in the public domain has increased, insights are most likely to come from its integration informatically in an open and accessible format. Here we present an online integrated immuno-analytic resource for the visualisation and extraction of SARS-CoV-2 meta-analysis data [10]. This platform utilises an automated pipeline for the formation of a whole genome sequence variant database for SARS-CoV-2 isolates worldwide (as of November 2020, n=150,090). We have integrated this dataset with a suite of B-cell epitope prediction platform meta analyses, HLA-I and HLA-II peptide prediction, an ‘epitope mapping’ analysis of available experimental *in vitro* confirmed epitope data from The Immune Epitope Database (IEDB) and a protein orthologue sequence analysis of six relevant coronavirus species (SARS, MERS, OC43, HKU1, NL63 and 229E); with all data updated and annotated regularly with information from the UniProt database. Furthermore, we have a feature that will enable users to visualise external analytical datasets presented in the literature (e.g. [11]). With this resource users can browse the SARS-CoV-2 proteome annotated with the above analyses and extract meta data to inform further experiments. To demonstrate the functionality of the immuno-analytics tool, we present an analysis of the SARS-CoV-2 spike, nucleocapsid and orf3a proteins, which are vaccine and serological targets.

## Methods

### Whole genome sequence data analysis

SARS-CoV-2 nucleotide sequences were downloaded from NCBI (https://www.ncbi.nlm.nih.gov) and GISAID (https://www.gisaid.org). As a part of an automated in-house pipeline, sequences were aligned using MAFFT software (v7.2) [12] and trimmed to the beginning of the first reading frame (orf1ab-nsp1). Sequences with >20% missing were excluded from the dataset. Using data available from the NCBI COVID-19 resource, a modified annotation (GFF) file was generated and open reading frames (ORFs) for each respective viral protein were extracted (taking in to account ribosomal slippage) using bedtools ‘getfasta’ function [13]. Each ORF was translated using EMBOSS transeq software [14] and the variants for each protein sequence were identified using an in-house script. As a part of our analysis pipeline we generated consensus sequences for each SARS-CoV-2 protein from the nucleotide database using the EMBOSS Cons CLI tool [14]. These canonical sequences were used as a reference for prediction, specificity and epitope mapping analyses.

### B-cell epitope prediction meta-analysis

Six epitope prediction software platforms were chosen for this analysis (Bepipred [15], AAPpred [16] DRREP [17], ABCpred [18], LBtope [19] and BCEpreds [20]). For each tool, we used the settings and quality cut-offs as recommended by their respective authors. The scores across the predictive platforms were then normalised (minimum-maximum scaled) to ensure that no single tool skewed the aggregate ‘consensus’ score, and combined to provide a single consensus B-cell epitope prediction score. Within the ‘raw data table’ (accessed from the tool’s landing page) users can dissect each score depending on their preference of methodology.

### HLA-I and HLA-II peptide prediction

We have incorporated an HLA-I peptide prediction analysis into the tool to aid in the scrutiny and development of vaccine candidates. CD8^+^effector immunity has been reported to play a central role in the response to SARS-CoV infection, as well as infection mediated immunopathology [21–23]. We used a database of 2,915 HLA-A, HLA-B and HLA-C alleles to make HLA-I peptide binding predictions using the netMHCpan server (v4.1) [24], with peptide lengths of 8 to 14 amino acids across the entire SARS-CoV-2 proteome. We chose to use the netMHCpan server for our HLA-I peptide prediction analysis, due to its high overall performance and its extensive HLA-I allele database [24]. We ran predictions for a total of 2,915 alleles (HLA-A 886, HLA-B 1412 and HLA-C 617), across all peptide lengths (8-14 amino acids). The analysis generated 1.1 billion candidates. After quality control we selected a total of 736,073 peptides based on strong binding affinity across the allele database for further analysis. We selected strong binding affinity peptides based on the tools internal binding scoring metrics. Only ‘strong binding’ alleles were selected for further analysis. For each position with a ligand with high binding affinity we analysed the percentage representation of the respective HLA-I type across the allele database. For predicting HLA-II peptides we used the MARIA online tool [25]. We pre-processed the SARS-CoV-2 canonical protein sequences using a 15 amino acid sliding window. Predictions were made for all available HLA-II alleles. A 95% cut off was chosen for a positive HLA-II presentation. All data for each 15-mer is displayed on the tool.

### Epitope mapping

B-cell epitopes for coronavirus species were sourced from the Immune Epitope Database (IEDB) resource (https://www.iedb.org). Using BLASTp [26] we mapped short amino acid epitope sequences onto the canonical sequence of SARS-CoV-2 proteins. A BLASTp bitscore of 25 with a minimum length of 8 residues was selected as a quality cut-off for mapped epitopes. The frequency of mapped epitopes was logged for each position in the protein and parsed for graphical representation.

### Coronavirus homology analysis

Reference proteomes for SARS, MERS, OC43, 229E, HKU1 and NL63 α and β coronavirus (-CoV) species were sourced from UniProt database. These sequences were processed into 10-mers using the *pyfasta* platform and mapped on to the canonical sequences of SARS-CoV-2 proteins using the aforementioned ‘epitope mapping’ process. The k-mer mapping technique applied used a matching threshold of at least 10 residues in orthologous viral proteins, which is of sufficient length to cover HLA-bound peptides and/or whole or part of a B-cell epitope; something that is challenging using only pairwise multiple sequence alignments. Homologous peptide sequences with a BLAST bitscore indicating 10 or more residues mapped to the target sequence were recorded and parsed for display on the graph.

### Online SARS-CoV-2 “Immuno-analytics” resource and analysis software

We developed an online immuno-analytics resource with an interactive plot that integrates SARS-CoV-2 genetic variation, epitope prediction and mapping, with other coronavirus homology, as well as a table for candidate proteome analysis. This tool is available online (from http://genomics.lshtm.ac.uk/immuno) (see **S1 Figure** for screenshots). The BioCircos.js library [10] was used to generate the interactive plot and *Datatables.net* libraries for the table. The underlying browser software and in-house pipelines for data analysis are available (https://github.com/dan-ward-bio/COVID-immunoanalytics). To facilitate the further scrutiny of non-synonymous mutations in the dataset, we have developed an interface as a part of the COVID immuno-analytics platform to enable users to temporally and geographically track mutant sites within a given SARS-CoV-2 protein Temporal and geographic data on individual mutations can be found on the ‘Mutation Tracker’ page, accessed via the tools home page. For the temporal/geographic non-synonymous mutation plots, we partitioned the whole genome sequencing dataset by week and continent and plotted non-synonymous allele frequencies using the Google Charts JavaScript libraries. Metadata on collection data and geographic location from each GISAID sequence was used to place each data point respectively. To improve sustainability of the tool, all functions and data of the website are generated and updated using automated data scripts developed in-house.

## Results

Analysis of 150,090 SARS-CoV-2 sequences identified 911,324 non-synonymous mutations across 16,951 sites in protein coding regions; 0.71% of these mutations are singleton events and 0.03% (46) of these mutations have a frequency above 1%, occurring in >1,500 samples. The most frequent mutations were the spike protein D614G (87.3%) and nsp12 L314P (87.5%), which were common across all the geographical regions (all >86%) **(Table 1)**, in keeping with their deep ancestral nature in the SARS-CoV-2 phylogenetic tree [23]. In particular, nsp12 L314P has been used to genotype the putative S and L strains of SARS-CoV-2, which have now been clustered into further groups [27]. Spike D614G lies 73 residues downstream from the spike RBD, a region of interest as it is a primary target of protective humoral responses and bears immunodominant epitopes that play a possible role in antibody dependant enhancement [28–31]. We have observed a strong correlation between the spatiotemporal accumulation of both spike D614G and nsp12 L314P **(Figure 1, Table 1),** due to either a common origin and subsequently linked accumulation by a founder effect, or a more complex biological interaction, including positive selection driven imparted by increased transmissibility as suggested by a recent study [32], Specifically, the spike D614G and nsp12 N314L both appear to have a near-identical frequency with a consistent increase across all geographic regions (negating weeks with poor data collection). In contrast, the frequency of orf3a Q57H appears to fluctuate, increasing and decreasing significantly from the time it was first observed in February 2020 (week 8) to present day (November 2020, week 43) (**Figure 1; Table 1**). Using the immuno-analytical tool, spike A222V, S477N and L18F variants were observed to have increased significantly in frequency between May and November 2020 (weeks 23-40). Spike mutations A222V and L18F appear to become entrenched in Europe reaching a total frequency of 70.6% and 31.6.%, respectively **(S1 Table, S2 Figure).** Moreover, A222V appears to be increasing in Asia and Oceania from week 41, and S477N has increased to >95% frequency across Oceania (N=8,321) with a peak of 9.3% in Europe **(S1 Table, S2 Figure),** consistent with a recent report [33].

**Figure 1.**
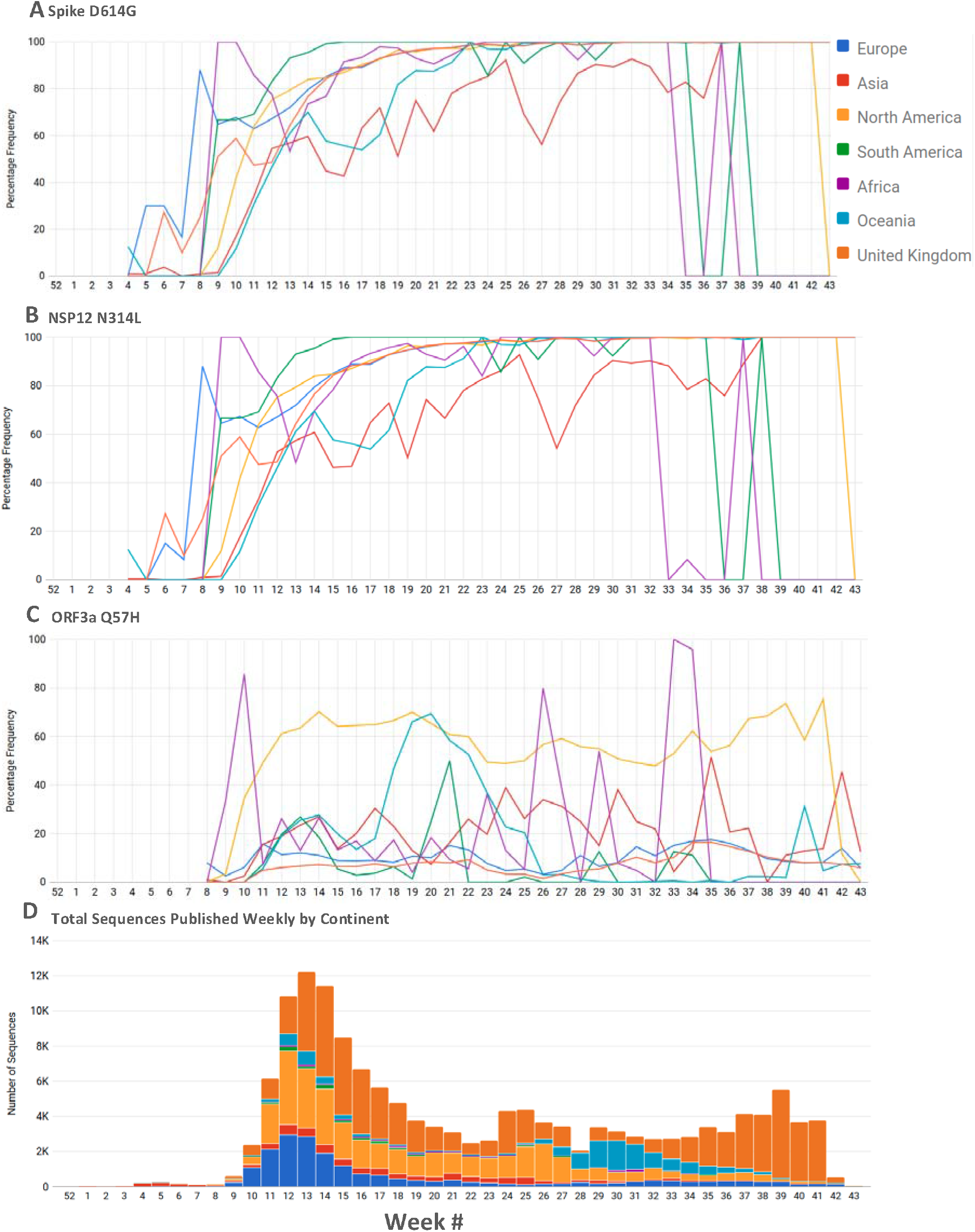
**(A-C)** High frequency non-synonymous spike D614G, NSP12 N314L and orf3A Q57H mutations found in SARS-CoV-2 plotted weekly by continent. This functionality is available on the website. Users can select any mutation from the main plot and visualise it temporally and geographically. UK sequences are not included in Europe due to high frequency. **(D)** A stacked bar chart representing total sequences published by each continent by week. This chart is included to assist users understand how allele frequencies may be affected by poor sampling. (http://genomics.lshtm.ac.uk/immuno/mutation_tracker).

**Table 1:** Most frequent non-synonymous mutations found in the 150,090 global SARS-CoV-2 whole genome sequences.

| Protein * | Pos. | Ref. Allele | Alt. Allele | Alt. Allele Freq. | Dominant Alt. Allele Freq. (150090) | Europe (88266) | Asia (7818) | NAm (38203) | SAm (1702) | AFR (1211) | OCE (12837) | UK (68017) ** |
| --- | --- | --- | --- | --- | --- | --- | --- | --- | --- | --- | --- | --- |
| **S** | 614 | D | V:G:S:N | 1:131490:2:10 | 0.877 | 0.975 | 0.940 | 0.866 | 0.890 | 0.866 | 0.868 | 0.883 |
| **nsp12** | 314 | N | H:F:L:S | 2:114:131053:1 | 0.876 | 0.973 | 0.938 | 0.863 | 0.887 | 0.876 | 0.865 | 0.881 |
| **N** | 203 | R | G:K:M:I:S | 9:57773:73:1:37 | 0.388 | 0.426 | 0.394 | 0.374 | 0.316 | 0.393 | 0.440 | 0.380 |
| **N** | 204 | G | R:V:L:Q:T | 57327:5:319:4:1 | 0.385 | 0.424 | 0.393 | 0.373 | 0.312 | 0.392 | 0.439 | 0.379 |
| **orf3a** | 57 | Q | K:Y:R:H:L | 1:6:6:35047:2 | 0.234 | 0.267 | 0.227 | 0.236 | 0.210 | 0.193 | 0.199 | 0.248 |
| **nsp2** | 85 | T | V:I | 1:25913 | 0.173 | 0.195 | 0.156 | 0.180 | 0.160 | 0.090 | 0.153 | 0.180 |
| **nsp6** | 37 | L | F | 11992 | 0.080 | 0.087 | 0.112 | 0.079 | 0.061 | 0.082 | 0.073 | 0.079 |
| **nsp2** | 120 | I | V:F:M | 7:11996:1 | 0.080 | 0.094 | 0.061 | 0.082 | 0.059 | 0.054 | 0.059 | 0.083 |
| **S** | 222 | A | T:V:P:I:S:F | 4:11819:2:1:12:1 | 0.079 | 0.099 | 0.148 | 0.057 | 0.177 | 0.135 | 0.026 | 0.085 |
| **orf10** | 30 | V | A:I:L | 2:2:11619 | 0.078 | 0.097 | 0.146 | 0.056 | 0.176 | 0.132 | 0.025 | 0.084 |
| **N** | 220 | A | V:T | 11555:5 | 0.077 | 0.097 | 0.146 | 0.056 | 0.176 | 0.131 | 0.025 | 0.083 |
| **S** | 477 | S | T:R:G:I:N:K | 1:19:2:59:9811:1 | 0.067 | 0.077 | 0.044 | 0.070 | 0.048 | 0.045 | 0.043 | 0.068 |
| **N** | 194 | S | A:L:P:T | 30:6441:2:1 | 0.043 | 0.052 | 0.043 | 0.035 | 0.031 | 0.053 | 0.038 | 0.049 |
| **orf8** | 84 | L | F:C:S:V | 1:1:6338:1 | 0.042 | 0.046 | 0.053 | 0.043 | 0.049 | 0.040 | 0.048 | 0.042 |
| **orf8** | 24 | S | L | 6151 | 0.041 | 0.053 | 0.033 | 0.036 | 0.034 | 0.009 | 0.022 | 0.048 |
| **S** | 18 | L | F:I | 5888:1 | 0.040 | 0.048 | 0.071 | 0.030 | 0.106 | 0.059 | 0.017 | 0.041 |
| **nsp5** | 15 | G | S:D | 5433:5 | 0.036 | 0.039 | 0.035 | 0.036 | 0.022 | 0.038 | 0.044 | 0.036 |
| **orf3a** | 251 | G | V:S:D:C | 5252:31:4:6 | 0.035 | 0.035 | 0.063 | 0.037 | 0.031 | 0.040 | 0.034 | 0.030 |
| **nsp13** | 541 | Y | C | 2606 | 0.017 | 0.019 | 0.032 | 0.017 | 0.018 | 0.021 | 0.019 | 0.017 |
| **nsp13** | 504 | P | L:H:S | 2535:1:111 | 0.017 | 0.020 | 0.032 | 0.017 | 0.018 | 0.022 | 0.021 | 0.017 |

Pos. = position; Freq. = Frequency; NAm = North America, SAm = South America, AFR = Africa, OCE =
Oceania; REF = Reference; ALT= Alternative; \* S = Spike, M = Membrane, N = Nucleocapsid, \*\* included
in Europe

The proximity of the D614G mutation to one of the functional domains of the spike protein has raised concerns, but whether it confers any gain in pathogenicity, transmissibility or immune evasion is still unclear [32]. Other high frequency mutations occur on the nucleocapsid gene (R203K, 38.8.0%; G204R 38.4%; across all geographical regions >31%; **Table 1**), which has been the target antigen for several serological RDTs currently in use or in production. Both of these mutations share a near-identical spaciotemporal profile. We have identified 363 non-synonymous variant sites across the nucleocapsid gene with mutations occurring 173,955 times in this dataset. Using the SARS-CoV-2 immuno-analytics platform we further queried these polymorphic regions for immunological relevance. The 20 residues surrounding the spike mutation D614G (S604-624) **(Figure 2)** have a high epitope prediction meta-score (34% increase on the global median) with 204 IEDB epitope positions mapping to the surrounding residues, suggesting this region is of high interest and may elicit a strong immune response. On top of the high level of SARS-CoV sequence homology reported, we have identified multiple clusters in the S2 domain of the spike protein, with homology to MERS, OC43, 229E, HKU1 and NL63 human coronaviruses, which may elicit a cross-reactive immune response in immune sera. Human coronavirus sequence homology is greatly reduced in the S1 domain, with only two small 10-residue pockets of OC43 and HKU1 identity (see **Figure 2).** We observed a 17% increase over the median epitope metascore within the receptor-binding motif (AA437-508), a region implicated in the direct ACE2 interaction. HLA-II peptide binding prediction (see **Figure 3)** yielded several epitopes within the receptor binding domain with high HLA-II ligand affinity, as well as strong B-cell epitope prediction scores (28% above the global median). Metadata obtained from the UniProt database reveals 3 clusters of glycosylated residues across the spike protein, a characteristic highlighted by this tool that should be considered when choosing expression systems for producing protein/peptides based on these regions.

**Figure 2.**
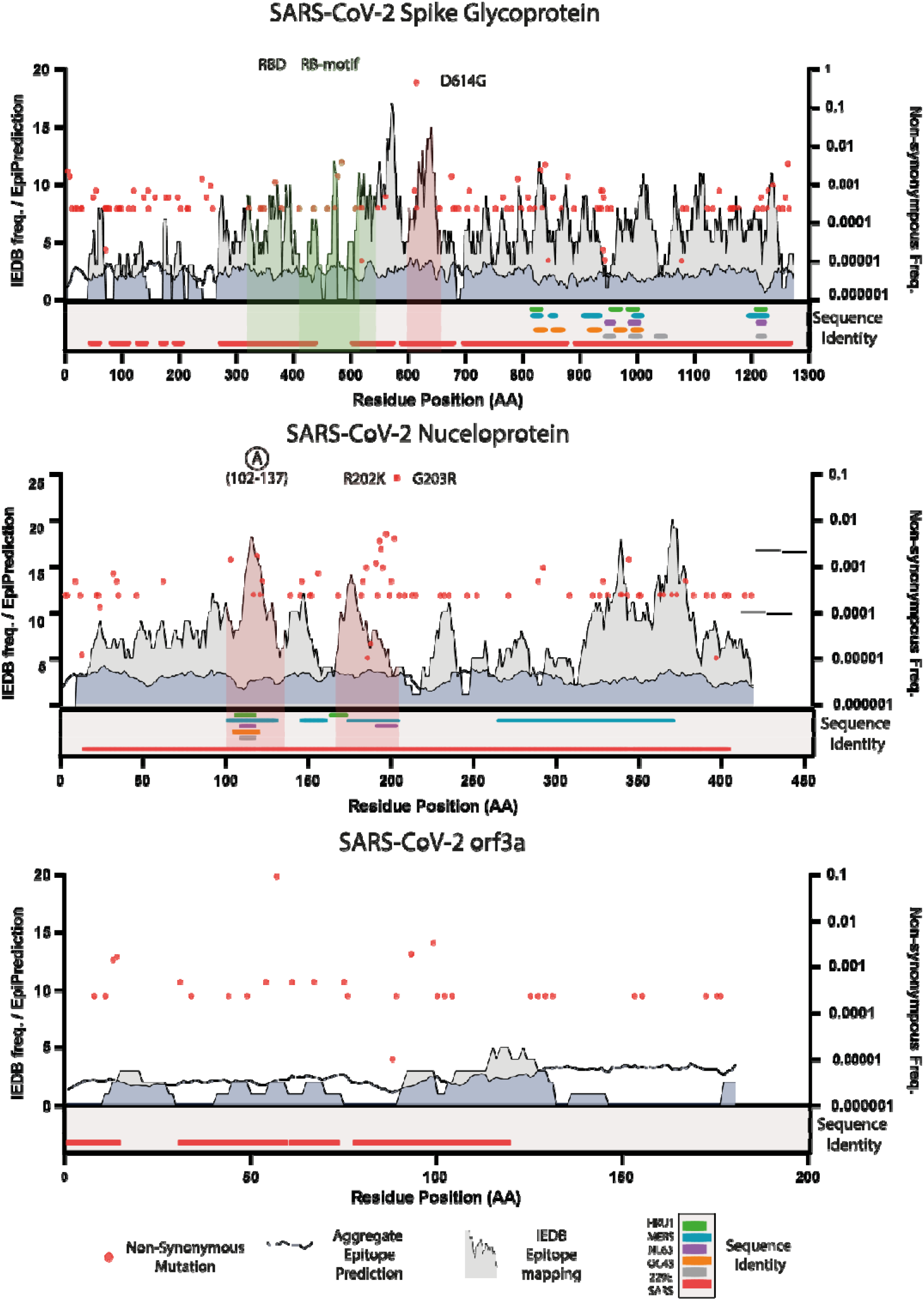
Linearised extracts from the SARS-CoV-2 immuno-analytics resource database; Spike, Nucleocapsid and orf3a proteins. Non-synonymous mutations, **B-cell** epitope prediction meta-score, IEDB epitope mapping and sequence identity analyses were plotted (see key). **Left axis** denotes scale for epitope prediction and IEDB mapping. **Right axis** denotes the relative allele frequency found in the genome dataset; AA amino acid.

**Figure 3.**
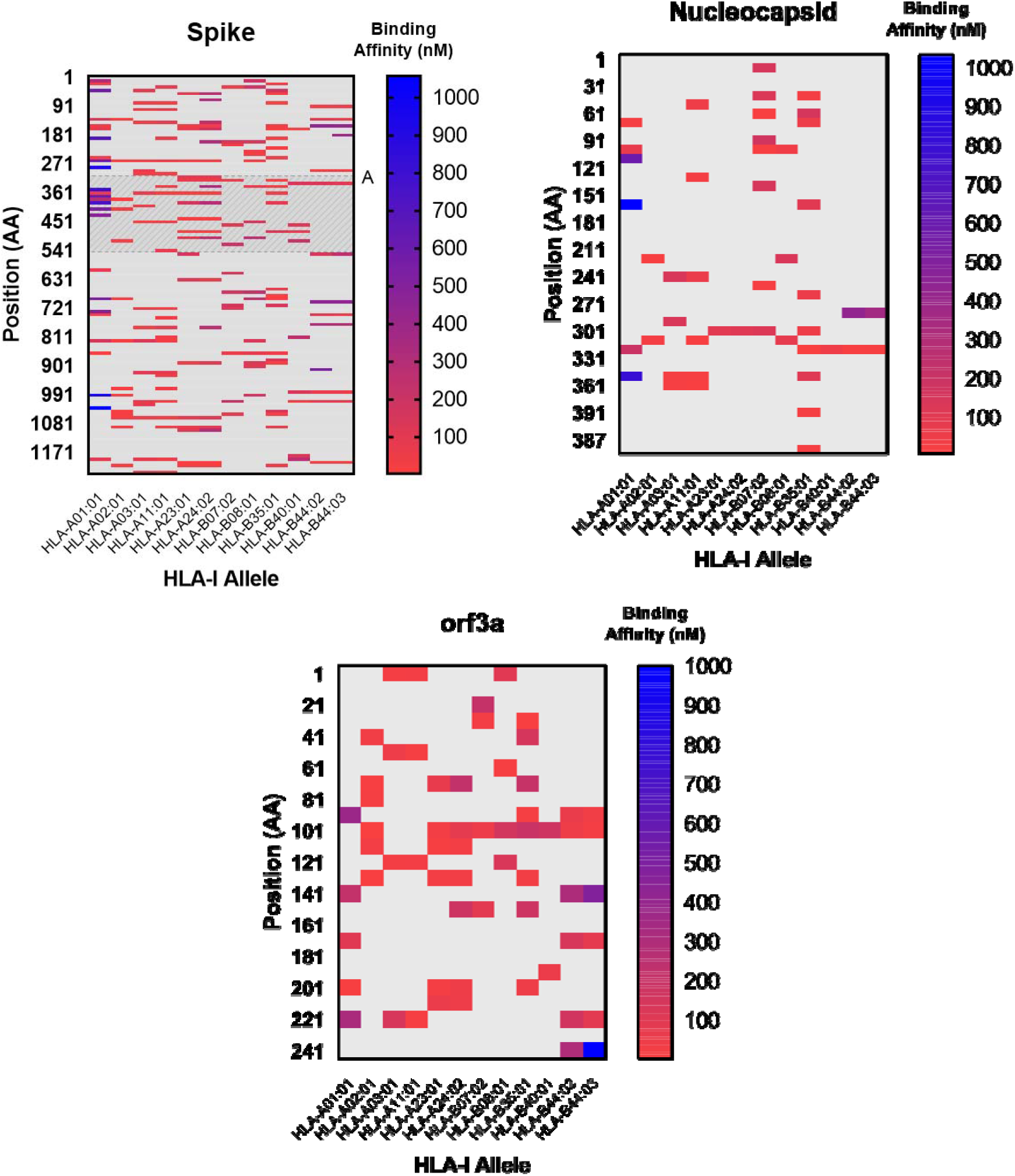
Mapping of HLA-I A and B epitopes to the spike, nucleocapsid and orf3a protein sequences. Epitopes were inferred using NetMHCpan 4.1. Only strong binding epitopes were selected for analysis, based on the tool’s internal classification system. A representative set of HLA-I alleles were used as described in Grifoni *et. al* (2020) [11]. Binding affinity is based on predicted IC50 values (nM) (a lower concentration equates to a stronger affinity). Shaded region (denoted by A) indicates the spike receptor binding domain (RBD).

For two high frequency non-synonymous nucleocapsid protein mutations (R203K and G204R; co-fixed), all but three of the 30 (N173-234) flanking residues have non-synonymous variants, with 5 sites reporting an alternative allele frequency greater than 1% (A220V, S194L, S197L, M2341 and P199L:S). The average epitope meta-score for these variant sites is 30% above the global median prediction score, with the two aforementioned high frequency mutant residues scoring 35% above the global median epitope predictive score. The sequence homology analysis of the nucleocapsid protein revealed a high level of shared identity between SARS-CoV (90%) and MERS-CoV (45%) on a per-residue basis. The nucleocapsid protein analysis revealed two clusters of shared human coronavirus orthologue identity **(Figure 2),** one of which was found to cross the aforementioned N173-234 region with identity to HKU1, NL63, MERS and SARS detected. Moreover, these clusters were found to have an increased IEDB epitope mapping frequency, high polymorphism frequency and B-cell epitope meta-scores (23% above the global median) indicative of potential B-cell immunogenicity. We focused on two nucleocapsid protein specific regions of interest (amino acids 102 to 137 and 167 to 206; **Figure 2).** Within the first 35-residue region (amino acids 102 to 137), we have detected NL63, SARS, OC43, 229E, MERS and HKU1 human coronavirus homology. Further, we observed an increase in mapped IEDB epitopes, including mapped linear peptidic B and T-cell epitopes from avian gammacoronavirus, murine betacoronavirus, feline and canine alphacoronavirus-1 providing *in vitro* confirmation that peptides within this region may indeed serve as immunogenic cross-reactive epitopes. The second region (amino acids 167 to 206) contains the R203K and G204R mutations along with a cluster of high frequency variants. We detected homology with HKU1, NL63 and MERS human coronavirus species along with a high frequency of SARS and murine coronavirus mapped IEDB epitopes, with a 34% increase on the median B-vell epitope prediction meta-score.

Previous studies of adaptive cellular effector immune responses to SARS-CoV infection have emphasised the importance of spike peptide presentation in the progression and severity of disease; regions of particular interest include: S436–443, S525–532, S366–374, S978, S1202 [21,22,34], We analysed these regions for their performance as HLA-I ligands *in-silico* and found that all of the regions of interest had a high binding affinity scores associated with that position. Moreover, these peptides were widely represented in the predictions made across the 2,915 HLA-A, −B and −C alleles used in this analysis. Taking into account all available HLA-A, B and C alleles, we found the spike peptides had an average allele coverage of 21%, 18% and 34% respectively. We performed an analysis to include 12 alleles with the highest frequency observed across the human population, as reported recently [11]. We found that peptides in the S366-374 and S1202 regions had high representation across the subset of 12 high frequency HLA-I alleles **(Figure 3).** These findings imply that the peptides as HLA-I ligands may have a putative role in initiating a protective cellular response in SARS-CoV-2 infections across a significant proportion of the HLA-I population worldwide. We have identified another region of interest that scores highly in the HLA-I peptide binding analysis. The S690-700 region has a high frequency of peptides with a high binding affinity with significant representation across all HLA-I alleles (HLA-A 40%, - B 23%, −C 60%). Furthermore, we have observed no mutations present in this region based on our SARS-CoV-2 variant analysis, implying this peptide appears to remain conserved making it a prime candidate for further study. The spike D614G mutation does not appear to have significantly elevated HLA-I epitope prediction scores **(Figure 3),** a finding supported by recent work [35]. The biological importance of spike D614G, particularly its immunological relevance and impact on transmission and disease, are still unclear [36,37].

Protein 3a (orf3a), has been reported to play a role in host immune modulation by decreasing interferon alpha-receptor expression in SARS-CoV infected cells and activating the NLRP3 inflammasome [38,39]; a response that may boost inflammation mediated COVID-19 pathology. Orf3a has been a target for SARS-CoV vaccinology studies, with reports of it eliciting potentially protective responses in both protein and DNA forms [23,40]. These immunogenic properties appear conserved in the SARS-CoV-2 orthologue, with consistently strong antibody responses reported in COVID-19 patients [41]. Looking across the SARS-CoV-2 proteome, of the 50 residues with the highest B-cell epitope prediction meta-score, orf3a occupies 16%, despite only constituting 2.5% of the total SARS-CoV-2 protein sequence. Moreover, there are numerous high affinity HLA-II epitopes, which may serve to elicit strong antibody responses. Although protein orf3a shares a high level of identity with its SARS-CoV orthologue, we detected no amino acid sequence homology with OC43, NL63, HKU1 and 229E human coronavirus species or any non-SARS-CoV IEDB epitopes.

Our analysis of the 150,090 SAR-CoV-2 whole genome sequences detected 267 variant sites within orf3a, with non-synonymous mutations occurring 68,473 times. A minority of these variant sites are singletons (8.2%) and five (1.8%) have a frequency higher than 1% (>1500 isolates), with a non-synonymous mutation density 40% lower than that of the nucleocapsid. The variant sites identified in the orf3a gene have a mean epitope predictive meta-score of 2.3, which is equal to the median global score, indicating that these sites may not form a part of a B-cell epitope. Comparing the predictive meta-scores of the nucleocapsid protein variant sites, we observed an increase of 26% over the global median, indicating that nucleocapsid protein non-synonymous mutations may impact epitope variability more than those found in orf3a. CD8^+^ effector responses to protein 3a have been characterised in SARS-CoV patients and appear to play a significant role in immunity [22,42,43]. Notably, alongside two within the spike protein, a peptide in orf3a (orf3a 36-50) has been found to form a part of the public (conserved) T-cell epitope repertoire across SARS-CoV patients [43]. This region scores highly in the HLA-II predictions with numerous HLA-A and HLA-B high affinity peptides covered (HLA-A 19%, −B 41%, −C 48%) and is relatively conserved with few low frequency non-synonymous mutations (maximum mutant allele frequency of 0.00219 (65 times)). We have identified one further region in orf3a (101-121) that scores highly with HLA-I epitope prediction across frequent HLA-I alleles **(Figure 3),** and therefore may be of interest to those studying HLA-I ligands. For HLA-I prediction, we observed that orf3a performs significantly better than the nucleocapsid. Despite the nucleocapsid protein sequence being 52% larger than that of orf3a, there are 34% more high affinity HLA-I epitopes across our subset of 12 frequent HLA-I alleles **(Figure 3),** which may indicate that orf3a has a more immunodominant role in cellular responses following intracellular processing when compared to the nucleocapsid protein.

## Discussion

We have developed an immuno-analytical tool that combines *in silico* prediction data with *in vitro* epitope mapping, SARS-CoV-2 genome variation and a k-mer-based human coronavirus sequence homology with curated functional annotation data. Furthermore, we have added functionality enabling users to track mutations geographically across time. An additional framework exists to annotate positions with relevant findings from the literature to further guide users’ research. The integration and co-visualisation of these data support the rational selection of diagnostics, vaccine targets with reverse-immunology, and highlight regions for further immunological studies. Using the tool, we demonstrate our analysis on three proteins and their mutant positions that are of relevance to current SARS-CoV-2 research, highlighting important features that will inform further SARS-CoV-2 research.

Understanding the magnitude of transmission and patterns of infection will lead to insights for postisolation strategies. The rapid emergence of the SARS-CoV-2 virus called for an expediated process to deploy serological RDTs for the detection of SARS-CoV-2 IgG/IgM antibody responses. There were reports early in the outbreak of lateral flow SARS-CoV-2 Ig RDTs not reaching sufficiently high levels of sensitivity and specificity [43]. While many assays use the spike protein as its sole antigen for antibody detection, others employ a combination of the spike and nucleocapsid proteins; other assays have been based solely on the nucleocapsid protein [6]. Our analyses suggest that, in its native form, the nucleocapsid protein may prove a sub-optimal target for use in serological diagnostic platforms. It possesses the greatest number of residues across all SARS-CoV-2 genes with high-frequency non-synonymous mutations, the majority of which have a high predictive epitope and IEDB epitope mapping scores when compared to variant positions of other genes. This implies that there may be an inherent variability in dominant antibody responses to different nucleocapsid protein isoforms, which may work to confound testing. We have located three regions of homology with other highly prevalent human coronavirus species, which could serve as non-specific SARS-CoV-2 epitopes if used in serological assays. Moreover, we have emphasised the high level of SARS-CoV identity across the SARS-CoV-2 proteome (except in orf8 and orf10), which may have implications for diagnostic deployment in countries that have had outbreaks involving SARS-CoV.

The spike protein has remained a focus of both vaccine and diagnostic research. Its functional role in viral entry imparts this antigen with immunodominant and neutralising antibody responses [28,44]. This role is reinforced in our analyses, with several clusters of high epitope meta-scores in functional regions, and IEDB epitope mapping counts. The S1 domain in particular has been the focus of a number of studies looking for specific antigens, not least because of its apparent lack of sequence homology with other human coronavirus species when compared to regions in the S2 domain, but also its strong functional and immunogenic role in SARS-CoV-2 infection [7,40,44,45]. However, as vaccination programmes begin, most of which will target the spike protein, it will become challenging to differentiate vaccination responses from those elicited by SARS-CoV-2 infection. There may then be a requirement for alternative viable targets for serological screening.

The broad nature of the analyses chosen for our tool may assist in the understanding of vaccine targets both during design and testing phases. The prediction of HLA-I ligands is relevant not only to the study of structural viral targets, but the full range of potentially immunologically relevant endogenous proteins analysed here that may be presented following intracellular processing, some of which may have less coverage in the literature. Our broad approach to HLA-I ligand prediction ensures that researchers understand the applicability of *in-silico* informed vaccine targets across different populations; a vital factor in pandemic situations. Further, ensuring that targets are both specific and devoid of polymorphism is essential to ensuring the longevity of vaccine responses and diagnostic capabilities, the analysis of which is achieved easily with our tool. The humoral and cellular immune responses, as well as the effects of H-CoV protein homology to SARS-CoV-2 proteins have yet to be fully characterised. With the significant levels of amino-acid sequence identity between SARS-CoV-2 and other H-CoV species detected in our analysis, researchers should be wary of the potentially deleterious effects of both non-specific humoral and cellular responses in enhancing infection, a phenomenon observed in a number of other viral pathology models. While the tracing and monitoring of non-synonymous mutations and their spatiotemporal analysis provides an initial indication of their importance, potentially the impact of evolutionary pressures on loci of interest, further analyses on signals of selection may provide additional insights. Computer intensive genome-wide analyses of positive selection are becoming available (e g. http://covid19.datamonkey.org/) and may be used to complement insights from our immuno-analytical tool.

Using the SARS-CoV-2 immuno-analytics platform we were able to identify shortcomings in current targets for diagnostics and suggest orf3a as another target for further study. This protein has proven *in vitro* immunogenicity in COVID-19 patients, as well as an array of supportive results from analyses performed here. The database underpinning the online tool is updated automatically using data parsing scripts that require minimal human curation. The monitoring of the temporal changes in the frequencies of mutations or their presence in multiple clades in a SARS-CoV-2 phylogenetic tree could provide insights for infection control, including post-vaccine introduction. Importantly, our open-access platform and tool enables the acquisition of all of the aforementioned data associated with the SARS-CoV-2 proteome, assisting further important research on COVID-19 control tools.

## Conclusions

The SARS-CoV-2 Immuno-analytics Platform enables the straightforward visualisation of multidimensional data to inform target selection in vaccine, diagnostic and immunology research. By integrating genomic and whole-proteome analyses with *in-silico* epitope predictions, we have highlighted important advantages and shortcomings of two proteins at the foci of COVID-19 research (spike and nucleocapsid), while suggesting another candidate for further study (orf3a). Both spike and nucleocapsid proteins have regions of high identity shared with other endemic H-CoV species. Moreover, several high frequency mutations found in our dataset lie within putative T and B-cell epitopes, something that should be taken into consideration when designing vaccines and diagnostics. Further, our work is likely to become more important as the introduction of vaccines will introduce new selection pressures that will need to be monitored for escape variations.

## DECLARATIONS

### Ethics approval and consent to participate

Not applicable

### Consent for publication

Not applicable

### Availability of data and materials

The sequencing data analysed during the current study are available from GISAID (https://www.gisaid.org) and NCBI (https://www.ncbi.nlm.nih.gov). Full analysis datasets can be downloaded from http://genomics.lshtm.ac.uk/immuno or https://github.com/dan-ward-bio/COVID-immunoanalytics.

### Authors’ contributions

DW, SC and TGC conceived and directed the project. MH and JEP provided software and informatic support. DW and JEP performed bioinformatic and statistical analyses under the supervision of TGC. DW, MLH, SC and TGC interpreted results. DW wrote the first draft of the manuscript. All authors commented and edited on various versions of the draft manuscript and approved the final manuscript. DW, TGC and SC compiled the final manuscript.

## Acknowledgements

We gratefully acknowledge the laboratories who submitted the data to the IEDB, NCBI and GISAID public databases on which this research is based. We also thank IEDB, NCBI and GISAID for developing and curating their databases. We gratefully acknowledge the availability of the Medical Research Council UK funded eMedLab (HDR UK) computing resource. DW wishes to thank Fraser, Josh, Stu and Paul for their support.

---

**SI Table.**
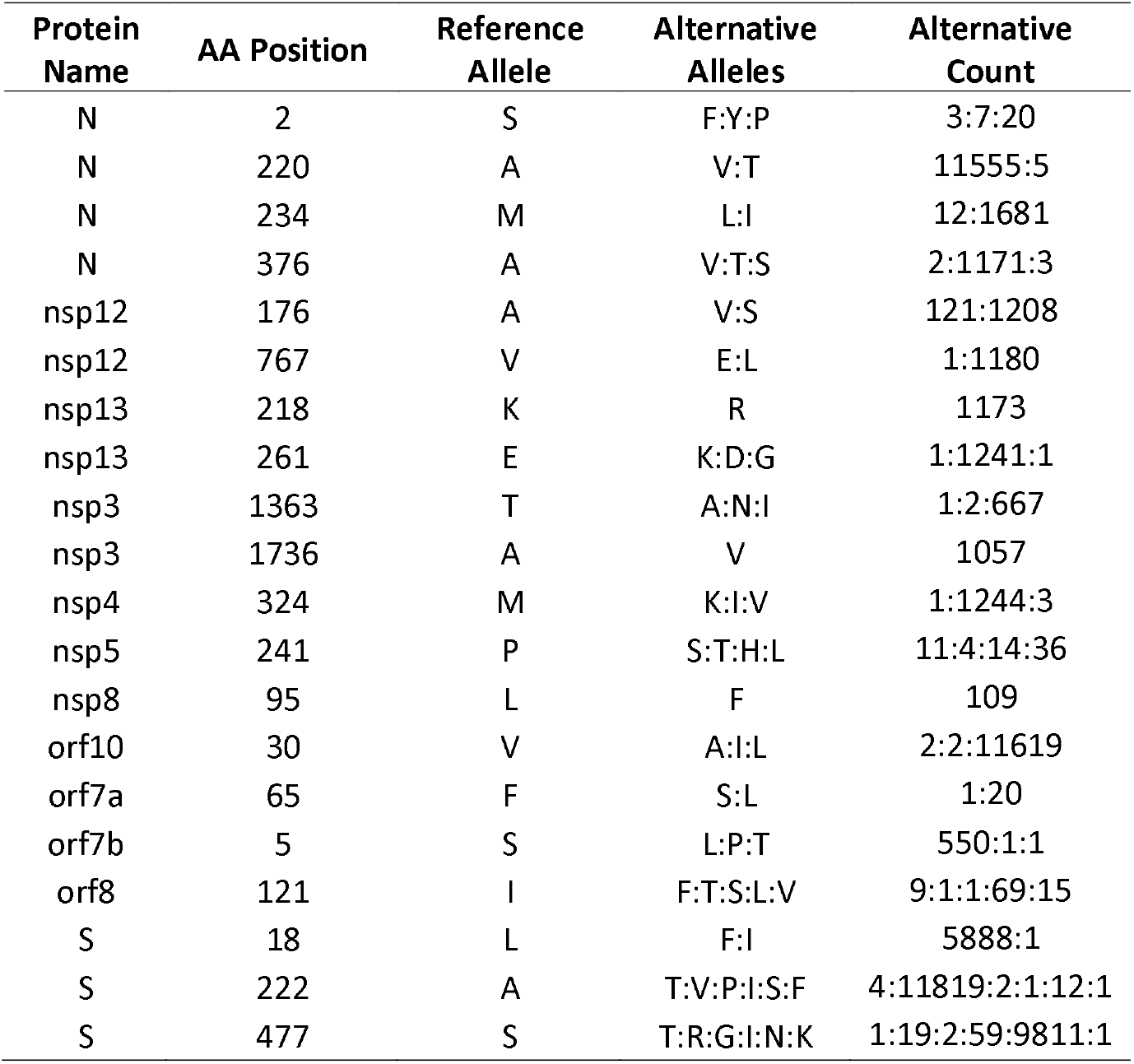
Extracted data from the Immuno-analytics tool. Mutations listed here have increased significantly over the 20-week period (weeks 20 to 40, year 2020) (see **S2 Figure** for spike mutations A222V, S477N and L18F).

**SI Figure.**
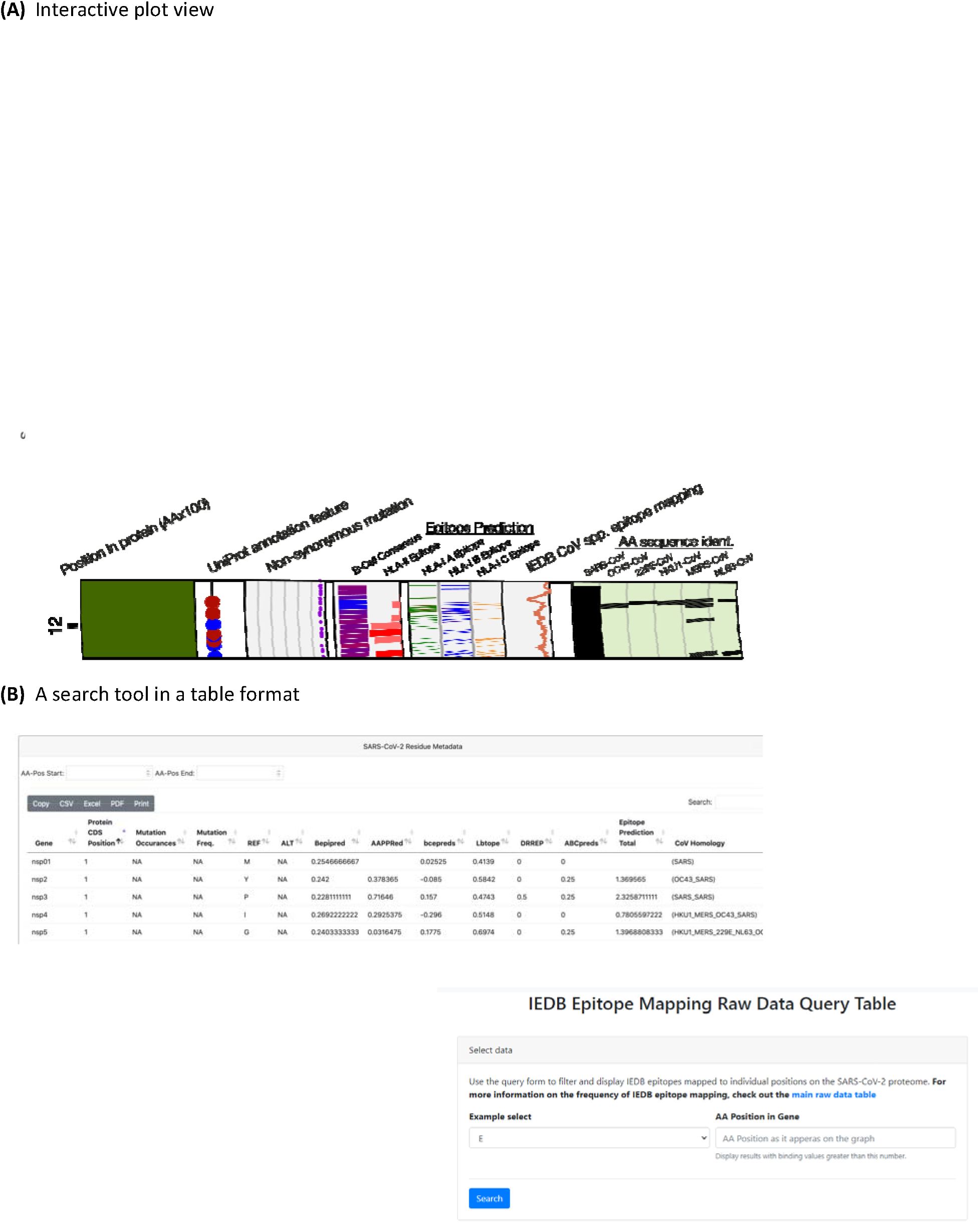
Screenshots from the Immuno-analytics webpage (http://genomics.lshtm.ac.uk/immuno). **(A)** Interactive plot view

**S2 Figure.**
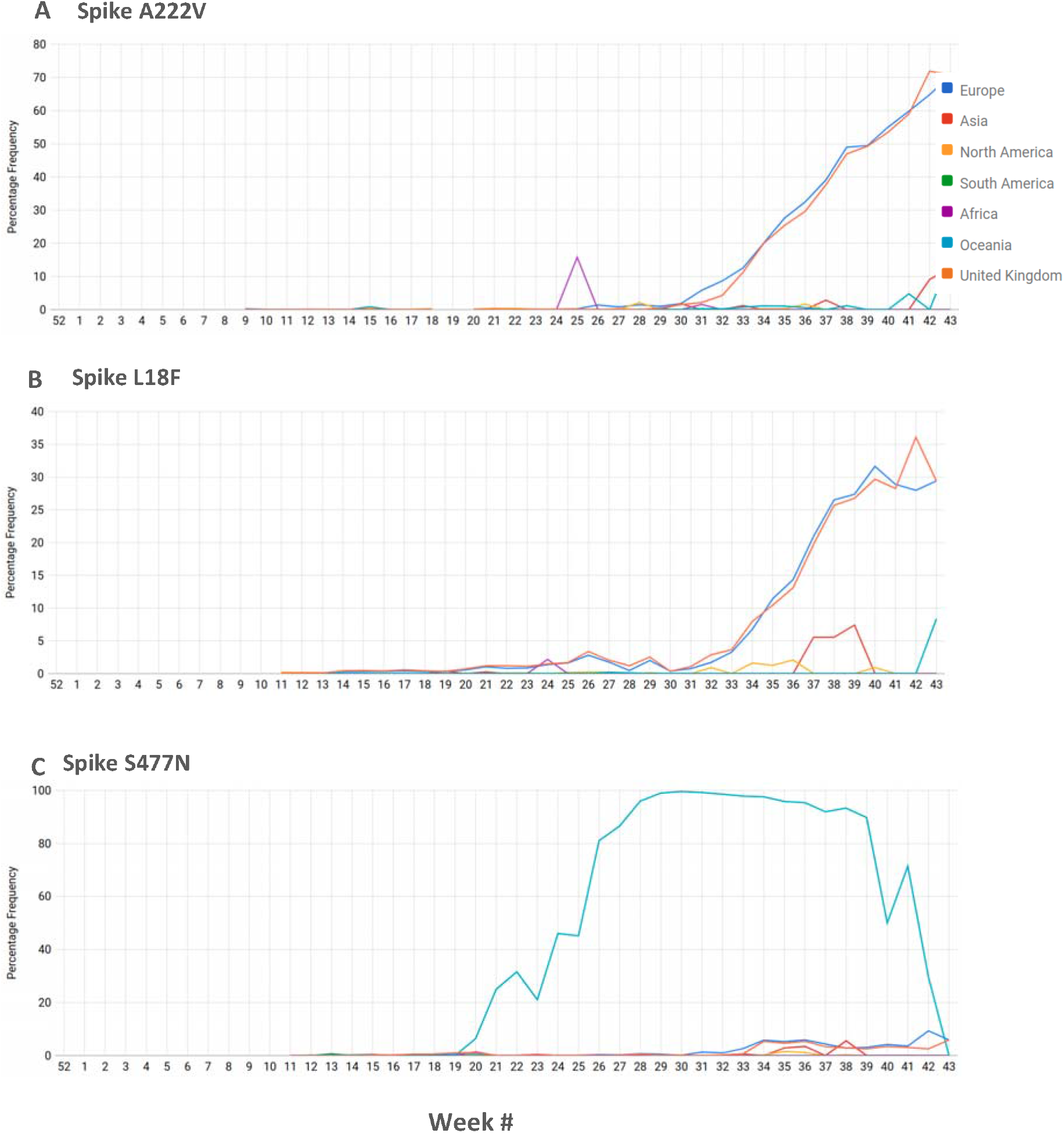
Screen capture from ‘Mutation Tracker’ page tracing spike mutations accumulating in Europe, North America and Oceania since the last week of December 2019 (week 52) into 2020 (week 1 onwards). Mutations can be traced across continents by week on the ‘Mutation Tracker’ page. Mutations shown here are in the Spike (A222V, S477N, and L18F).

